## Supplemental Figures and Tables for "An interbacterial DNA deaminase toxin directly mutagenizes surviving target populations"

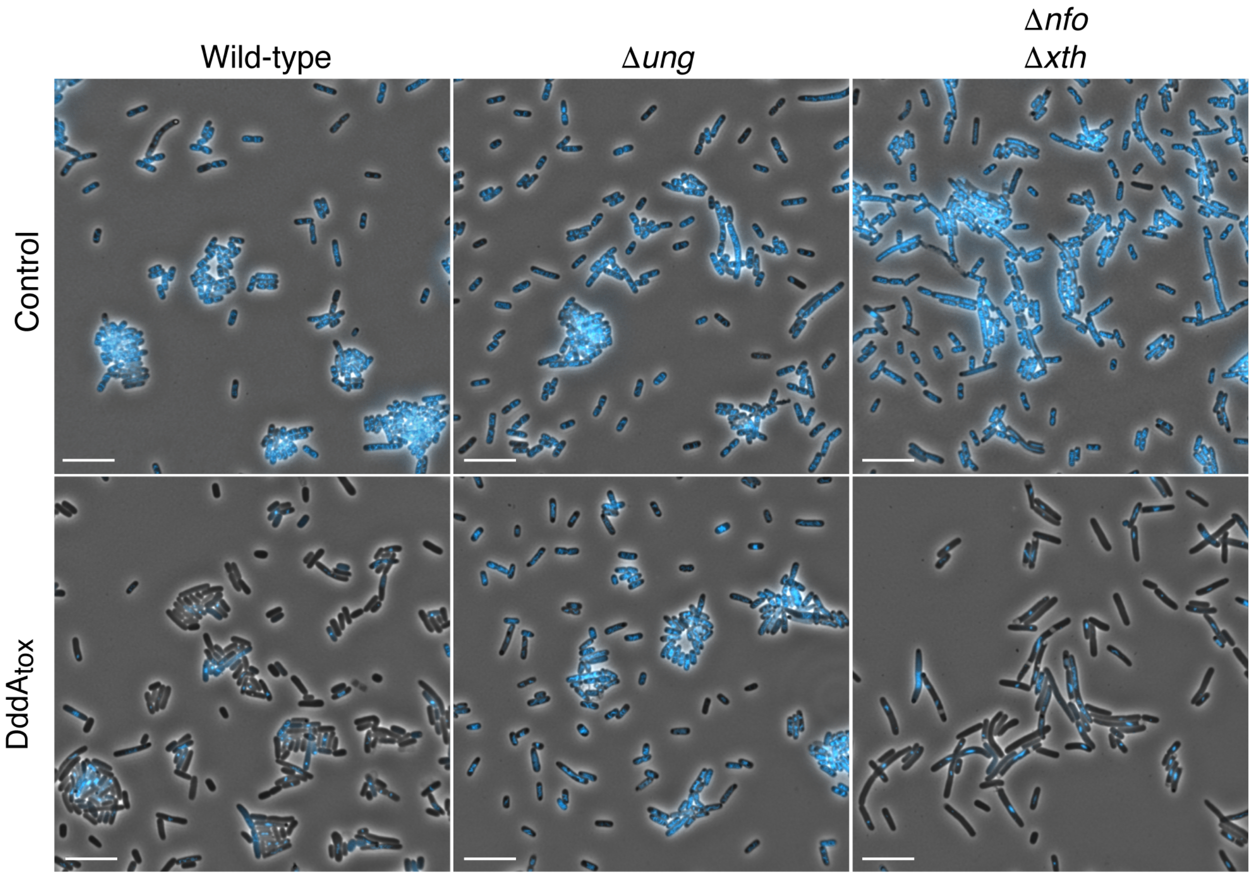

**Supplemental Figure 1. Expression of DddA$_{tox}$ leads to nucleoid degradation in *E. coli***. Full field view of fluorescence micrograph shown in Figure 1 (A) depicting *E. coli* strains expressing DddA$_{tox}$ or carrying an empty vector (Control). DAPI staining (DNA) is shown in cyan. Scale bar = 10μM.

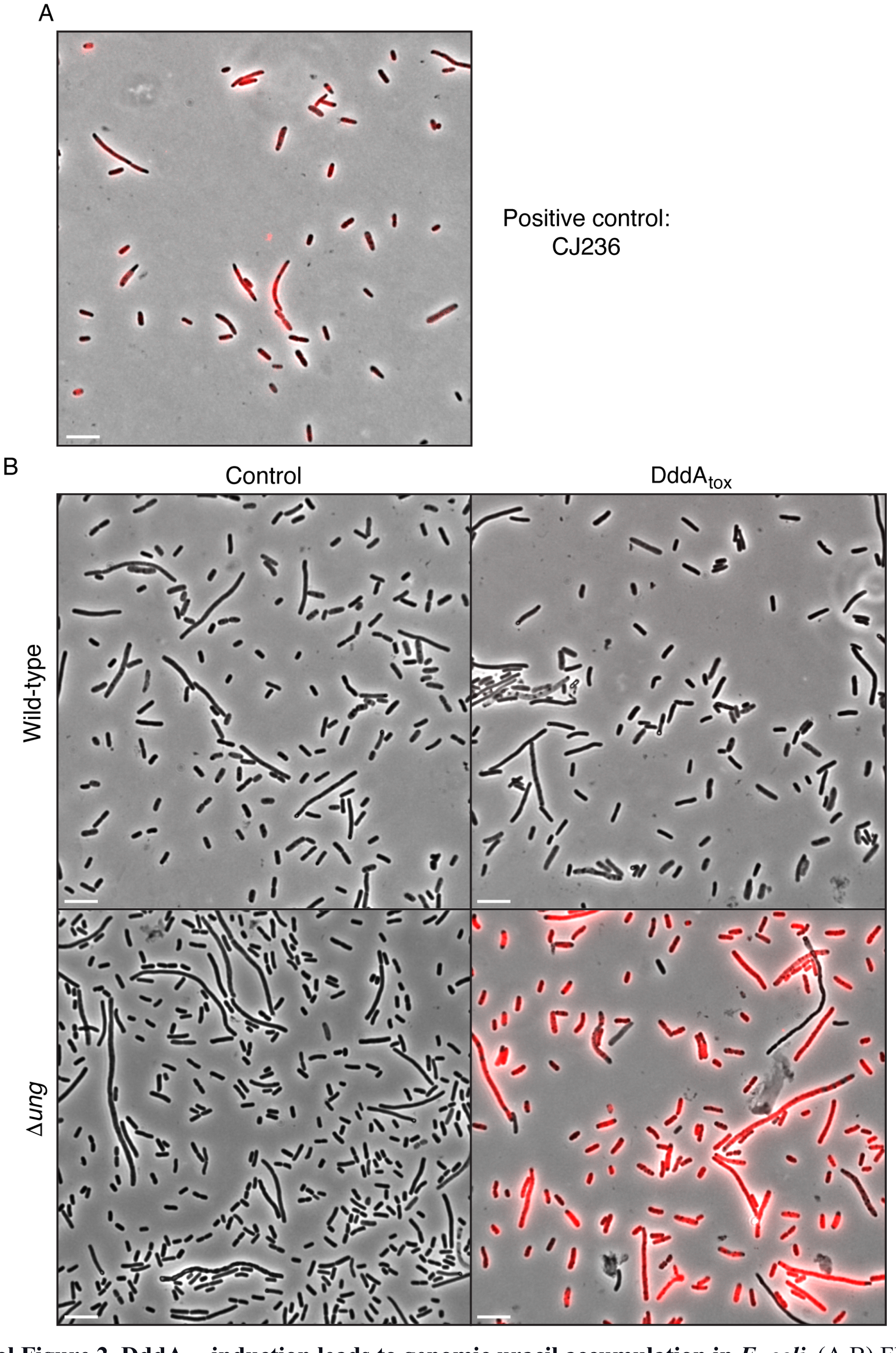

**Supplemental Figure 2. DddA$_{tox}$ induction leads to genomic uracil accumulation in *E. coli*.** (A,B) Fluorescence microscopy indicating genomic uracil incorporation (red) in the noted *E. coli* strains. Scale bar = 10μM. (A) Staining of a positive control strain for uracil incorporation, *E. coli* CJ236 [*dut−, ung−*]. This strain accumulates ~500-fold more uracil than wild-type. (B) Complete fields of view used to generate fluorescence micrographs depicted in Figure 1D.

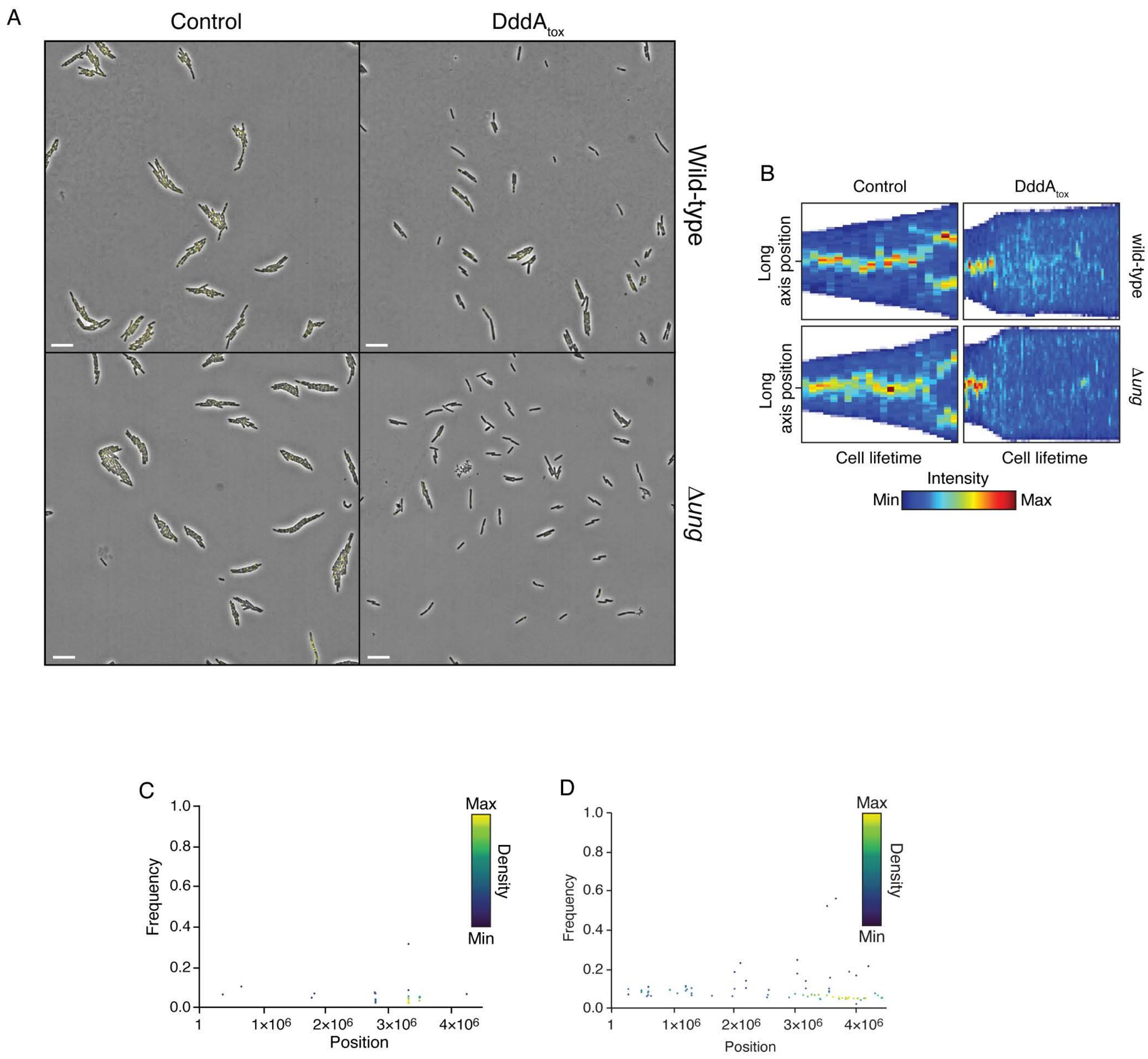

**Supplemental Figure 3. Expression of DddA$_{tox}$ in *E. coli* leads to replication arrest but does not mutagenize RNA.** (A) Full field of view of fluorescence micrographs shown in Figure 2B. Scale bar = 10μM. (B) Kymograph of representative cells from time-lapse microscopy of DnaN-YPet expressing strains during a 6 hr timelapse post-induction of DddA$_{tox}$ or empty vector (Control), scale bar = 5 μm. (C-D) Representation by chromosomal position, frequency, and density of SNVs detected with RNA-seq analysis of total cDNA collected from *E. coli* 60 min after induction of DddA$_{tox}$ (C) or the empty vector (Control) (D).

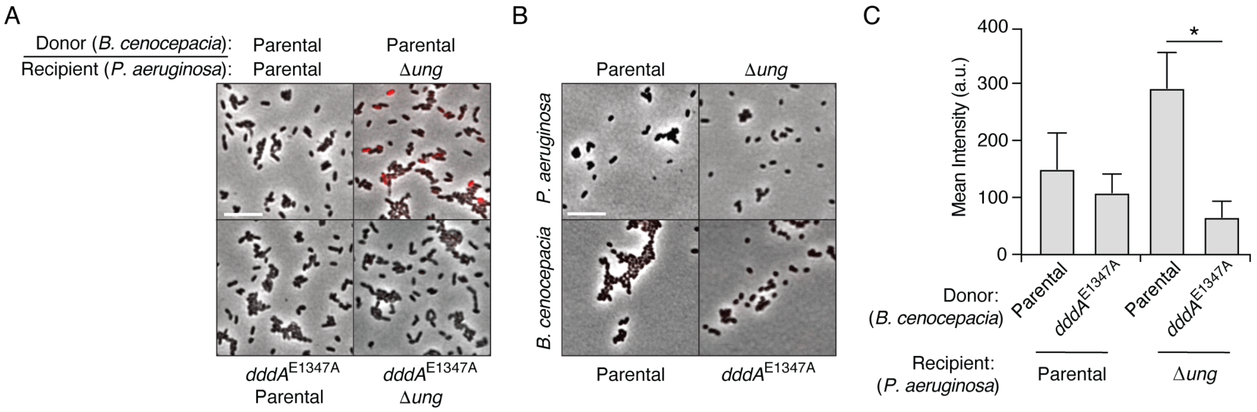

**Supplemental Figure 4. *P. aeruginosa* accumulates genomic uracil during competition with *B. cenocepacia*.** (A-B) Fluorescence microscopy indicating uracil incorporation (red) into genomic DNA of cells recovered after one hour of growth from cocultures of *P. aeruginosa* wild-type or Δ*ung* cells and the indicated strains of *B. cenocepacia* (A) or each of the strains used in (A) grown in monoculture (B). (C) Quantification of uracil labeling signal from cells shown in A (n = ~50 cells per condition). Values and error bars reflect mean ± s.d. of n = 2 independent biological replicates. *P < 0.0001

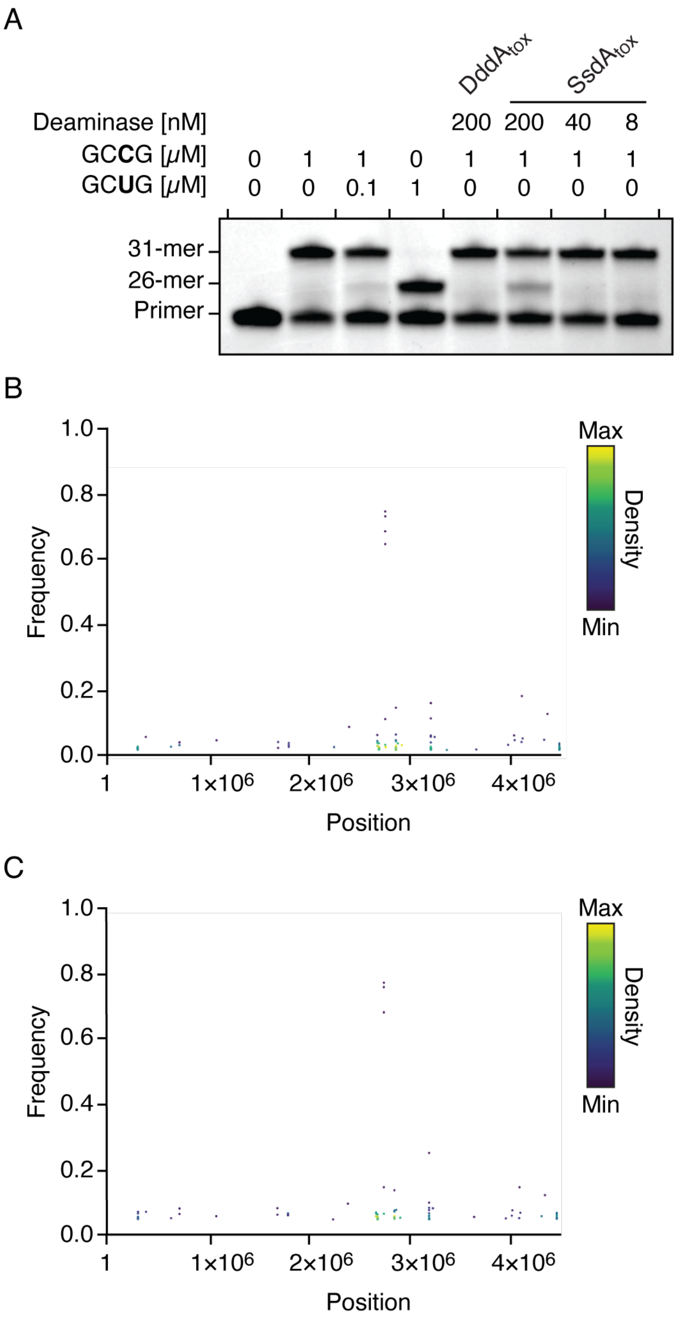

**Supplemental Figure 5. SsdA$_{tox}$ exhibits cytidine deaminase activity toward RNA *in vitro*, but not *in vivo*.** (A) Poisoned primer extension assay to detect deamination of cytidine in single-stranded RNA substrates. Deamination activity generates a 26-mer product. Images are representative of n = 2 independent replicates. (B-C) Representation of SNVs by chromosomal position, frequency, and density detected with RNA-seq analysis of total cDNA collected from *E. coli* 60 min after induction of SsdA$_{tox}$ (B) or an empty vector (Control) (C).

**Supplemental Video 1. DddA inhibits *E. coli* growth.** Time lapse phase microscopy of *E. coli* expressing DddA during a 300 min period.

**Supplemental Video 2. DddA arrests replication in *E. coli* growth.** Time lapse fluorescence microscopy of *E. coli* with YPet-labeled DnaN expressing DddA during a 300 min period

**Supp. Table 1. Summary of SNPs in cells obtained from coculture with *B. cenocepacia*.**

| ***E. coli* AB1157 NC_000913.3 WGS** | | | |
|---|---|---|---|
| **Isolate** | **Position** | **Context** | **DddA signature (Yes/No)** |
| AB1157-A2 | 2,573,448 | G**G**A | Y |
| AB1157-A6 | 636,682 | A**G**A | Y |
| AB1157-A7 | 561,441 | G**G**A | Y |
| AB1157-A7 | 2,380,964 | T**C**C | Y |
| AB1157-A7 | 4,546,218 | T**C**G | Y |
| AB1157-A8 | 700,660 | C**G**A | Y |
| AB1157-A8 | 903,356 | T**C**C | Y |
| AB1157-A8 | 1,019,069 | C**G**A | Y |
| AB1157-A8 | 2,180,410 | T**C**C | Y |
| AB1157-A8 | 4,023,769 | C**G**A | Y |
| AB1157-B3 | 457,974 | T**C**C | Y |
| AB1157-B4 | 682,518 | **TG**A | Y |

| ***E. coli* AB1157 NC_000913.3 *rpoB*** | | | |
|---|---|---|---|
| **Isolate** | **Position** | **Context** | **DddA signature (Yes/No)** |
| AB1157-A2 | 4,182,779 | T**C**T | Y |
| AB1157-A4 | 4,182,790 | G**G**A | Y |
| AB1157-A6 | 4,182,779 | T**C**T | Y |
| AB1157-A7 | 4,182,779 | T**C**T | Y |
| AB1157-A8 | 4,182,779 | T**C**T | Y |
| AB1157-B3 | 4,182,779 | T**C**T | Y |
| AB1157-B4 | 4,182,779 | T**C**T | Y |
| AB1157-C1 | 4,182,779 | T**C**T | Y |
| AB1157-C10 | 4,182,779 | T**C**T | Y |
| AB1157-C11 | 4,182,779 | T**C**T | Y |
| AB1157-C12 | 4,182,790 | G**G**A | Y |
| AB1157-C2 | 4,182,779 | T**C**T | Y |
| AB1157-C3 | 4,182,779 | T**C**T | Y |
| AB1157-C4 | 4,182,779 | T**C**T | Y |
| AB1157-C5 | 4,182,779 | T**C**T | Y |
| AB1157-C6 | 4,182,836 | T**C**C | Y |
| AB1157-C7 | 4,182,779 | T**C**T | Y |
| AB1157-C8 | 4,182,779 | T**C**T | Y |
| AB1157-C9 | 4,182,779 | T**C**T | Y |
| AB1157-D1 | 4,182,779 | T**C**T | Y |
| AB1157-D2 | 4,182,779 | T**C**T | Y |
| AB1157-D3 | 4,182,779 | T**C**T | Y |
| AB1157-D4 | 4,182,779 | T**C**T | Y |
| AB1157-D5 | 4,182,779 | T**C**T | Y |

| ***E. coli* EHEC NZ_CP008957.1 WGS** |
|---|

| Isolate | Position | Context | DddA signature (Yes/No) |
|---|---|---|---|
| EHEC-A2 | 745,731 | T**C**G | Y |
| EHEC-A2 | 839,041 | C**G**A | Y |
| EHEC-A2 | 2,166,407 | C**G**A | Y |
| EHEC-A2 | 4,797,858 | T**C**G | Y |
| EHEC-A2 | 5,079,358 | T**C**C | Y |
| EHEC-A2 | 5,276,086 | C**G**A | Y |
| EHEC-A3 | 1,614,650 | T**C**G | Y |
| EHEC-A3 | 2,231,975 | T**C**G | Y |
| EHEC-A3 | 2,523,951 | C**G**A | Y |
| EHEC-A3 | 3,267,194 | C**G**A | Y |
| EHEC-A3 | 3,370,445 | T**C**G | Y |
| EHEC-A3 | 3,444,101 | T**C**G | Y |
| EHEC-A3 | 3,928,392 | T**C**G | Y |
| EHEC-A3 | 5,079,358 | T**C**C | Y |
| EHEC-A4 | 1,330,651 | G**G**A | Y |
| EHEC-A4 | 2,175,988 | C**G**A | Y |
| EHEC-A4 | 4,851,817 | T**C**G | Y |
| EHEC-A4 | 5,079,358 | T**C**C | Y |
| EHEC-A4 | 5,466,257 | T**C**G | Y |
| EHEC-B1 | 1,242,341 | T**C**G | Y |
| EHEC-B1 | 3,849,029 | T**C**G | Y |
| EHEC-B1 | 5,079,358 | T**C**C | Y |
| EHEC-B1 | 5,428,118 | C**G**A | Y |
| EHEC-B2 | 3,706,386 | C**G**A | Y |
| EHEC-B2 | 4,623,007 | C**G**A | Y |
| EHEC-B2 | 5,079,358 | T**C**C | Y |
| EHEC-B3 | 784,298 | T**C**C | Y |
| EHEC-B3 | 920,896 | T**C**G | Y |
| EHEC-B3 | 2,142,210 | C**G**A | Y |
| EHEC-B3 | 3,144,163 | T**C**G | Y |
| EHEC-B3 | 3,190,034 | T**C**G | Y |
| EHEC-B3 | 3,874,627 | C**G**A | Y |
| EHEC-B3 | 5,079,358 | T**C**C | Y |
| EHEC-B3 | 5,280,427 | A**G**A | Y |
| EHEC-B4 | 10,950 | T**C**G | Y |
| EHEC-B4 | 195,295 | T**C**G | Y |
| EHEC-B4 | 817,797 | T**C**G | Y |
| EHEC-B4 | 1,009,128 | T**C**C | Y |
| EHEC-B4 | 1,199,226 | T**C**G | Y |
| EHEC-B4 | 2,381,467 | C**G**A | Y |
| EHEC-B4 | 3,121,763 | T**C**G | Y |
| EHEC-B4 | 3,288,109 | T**C**C | Y |
| EHEC-B4 | 3,313,845 | T**C**G | Y |
| EHEC-B4 | 3,443,031 | T**C**C | Y |

| | | | |
|---|---|---|---|
| EHEC-B4 | 3,603,862 | T**C**C | Y |
| EHEC-B4 | 3,613,539 | C**G**A | Y |
| EHEC-B4 | 3,855,072 | G**G**A | Y |
| EHEC-B4 | 4,573,974 | C**G**A | Y |
| EHEC-B4 | 4,948,346 | T**C**C | Y |
| EHEC-B4 | 5,104,176 | G**G**A | Y |
| EHEC-B4 | 5,525,915 | C**G**A | Y |
| EHEC-B4 | 13,936 | G**C**G | N |

| ***E. coli* EHEC NZ_CP008957.1 *rpoB*** | | | |
|---|---|---|---|
| **Isolate** | **Position** | **Context** | **DddA signature (Yes/No)** |
| EHEC-A2 | 4,771,622 | G**G**A | Y |
| EHEC-A3 | 4,771,622 | G**G**A | Y |
| EHEC-A4 | 4,771,622 | G**G**A | Y |
| EHEC-B1 | 4,771,622 | G**G**A | Y |
| EHEC-B2 | 4,771,622 | G**G**A | Y |
| EHEC-B3 | 4,771,622 | G**G**A | Y |
| EHEC-B4 | 4,771,622 | G**G**A | Y |
| EHEC-C1 | 4,771,622 | G**G**A | Y |
| EHEC-C2 | 4,771,622 | G**G**A | N |
| EHEC-C3 | 4,771,622 | G**G**A | N |
| EHEC-C4 | 4,771,622 | G**G**A | Y |
| EHEC-C5 | 4,771,622 | G**G**A | Y |
| EHEC-C6 | 4,771,622 | G**G**A | N |
| EHEC-C7 | 4,771,668 | T**C**C | Y |

| ***K. pneumoniae* CP000647.1 WGS** | | | |
|---|---|---|---|
| **Isolate** | **Position** | **Context** | **DddA signature (Yes/No)** |
| KP-A1 | 268,966 | T**C**G | Y |
| KP-A1 | 630,584 | C**G**A | Y |
| KP-A1 | 972,660 | T**C**A | Y |
| KP-A1 | 1,056,047 | T**C**G | Y |
| KP-A1 | 1,111,902 | C**G**A | Y |
| KP-A1 | 1,188,364 | T**C**G | Y |
| KP-A1 | 1,933,442 | T**C**G | Y |
| KP-A1 | 2,664,710 | C**G**A | Y |
| KP-A1 | 3,098,987 | T**C**G | Y |
| KP-A1 | 3,815,785 | T**C**G | Y |
| KP-A1 | 4,237,588 | T**C**G | Y |
| KP-A1 | 4,343,291 | C**G**A | Y |
| KP-A1 | 4,416,303 | C**G**A | Y |
| KP-A2 | 400,833 | T**C**G | Y |
| KP-A2 | 441,001 | C**G**A | Y |
| KP-A2 | 443,547 | T**C**G | Y |

| | | | |
|---|---|---|---|
| KP-A2 | 596,586 | T**C**G | Y |
| KP-A2 | 699,516 | T**C**G | Y |
| KP-A2 | 1,144,391 | T**C**G | Y |
| KP-A2 | 1,374,149 | T**C**G | Y |
| KP-A2 | 1,646,149 | C**G**A | Y |
| KP-A2 | 1,765,683 | T**C**G | Y |
| KP-A2 | 1,875,143 | C**G**A | Y |
| KP-A2 | 1,953,329 | C**G**A | Y |
| KP-A2 | 2,626,210 | C**G**A | Y |
| KP-A2 | 2,695,891 | C**G**A | Y |
| KP-A2 | 3,304,887 | T**C**G | Y |
| KP-A2 | 3,718,513 | T**C**G | Y |
| KP-A2 | 3,816,428 | T**C**G | Y |
| KP-A2 | 4,267,282 | T**C**G | Y |
| KP-A2 | 4,333,019 | C**G**A | Y |
| KP-A2 | 4,649,355 | T**C**G | Y |
| KP-A2 | 4,674,778 | C**G**A | Y |
| KP-A2 | 4,857,315 | C**G**A | Y |
| KP-A2 | 5,305,473 | T**C**G | Y |
| KP-A3 | 132,760 | T**C**G | Y |
| KP-A3 | 782,378 | T**C**G | Y |
| KP-A3 | 1,048,665 | T**C**G | Y |
| KP-A3 | 1,406,220 | T**C**G | Y |
| KP-A3 | 1,406,242 | C**G**A | Y |
| KP-A3 | 1,542,047 | C**G**A | Y |
| KP-A3 | 1,930,481 | G**C**C | Y |
| KP-A3 | 2,121,913 | T**C**G | Y |
| KP-A3 | 2,211,040 | C**G**A | Y |
| KP-A3 | 2,228,986 | T**C**G | Y |
| KP-A3 | 2,620,049 | T**C**G | Y |
| KP-A3 | 2,862,127 | T**C**G | Y |
| KP-A3 | 3,239,414 | C**G**A | Y |
| KP-A3 | 3,722,729 | T**C**G | Y |
| KP-A3 | 5,070,069 | C**G**A | Y |
| KP-A3 | 5,089,002 | G**G**A | Y |
| KP-A3 | 5,172,472 | T**C**G | Y |
| KP-A3 | 5,245,737 | T**C**G | Y |
| KP-B1 | 453,408 | C**G**A | Y |
| KP-B1 | 766,528 | C**G**A | Y |
| KP-B1 | 949,753 | C**G**G | N |
| KP-B1 | 2,254,942 | T**C**G | Y |
| KP-B1 | 2,318,894 | T**C**G | Y |
| KP-B1 | 2,637,023 | C**G**A | Y |
| KP-B1 | 3,063,632 | C**G**A | Y |
| KP-B1 | 3,133,040 | T**C**C | Y |

| | | | |
|---|---|---|---|
| KP-B1 | 3,345,186 | T**C**G | Y |
| KP-B1 | 3,519,995 | T**C**G | Y |
| KP-B1 | 4,046,644 | T**C**G | Y |
| KP-B1 | 4,600,477 | T**C**G | Y |
| KP-B1 | 4,651,523 | G**C**G | N |
| KP-B1 | 5,313,678 | T**C**G | N |
| KP-B2 | 327,129 | T**C**G | Y |
| KP-B2 | 1,400,074 | C**C**G | N |
| KP-B2 | 2,254,942 | T**C**G | Y |
| KP-B2 | 2,318,894 | T**C**G | Y |
| KP-B2 | 3,049,130 | T**C**G | Y |
| KP-B2 | 3,063,632 | C**G**A | Y |
| KP-B2 | 3,133,040 | T**C**C | Y |
| KP-B2 | 3,519,995 | T**C**G | Y |
| KP-B2 | 4,046,644 | T**C**G | Y |
| KP-B2 | 5,313,678 | T**C**G | N |
| KP-B3 | 923,447 | T**C**G | Y |
| KP-B3 | 1,400,074 | C**C**G | N |
| KP-B3 | 1,725,719 | T**C**C | N |
| KP-B3 | 2,354,429 | C**G**A | Y |
| KP-B3 | 2,664,710 | C**G**A | Y |
| KP-B3 | 3,014,427 | C**G**A | Y |
| KP-B3 | 3,345,186 | T**C**G | Y |
| KP-B3 | 3,851,167 | T**C**G | Y |
| KP-B4 | 327,129 | T**C**G | Y |
| KP-B4 | 1,400,074 | C**C**G | N |
| KP-B4 | 2,254,942 | T**C**G | Y |
| KP-B4 | 2,318,894 | T**C**G | Y |
| KP-B4 | 2,926,868 | G**G**A | Y |
| KP-B4 | 2,956,330 | C**G**A | Y |
| KP-B4 | 3,345,186 | T**C**G | Y |
| KP-B4 | 3,786,107 | T**C**G | Y |
| KP-B4 | 4,055,568 | T**C**G | Y |
| KP-B4 | 5,313,678 | T**C**G | N |
| KP-B5 | 1,386,432 | C**G**A | Y |
| KP-B5 | 1,400,074 | C**C**G | N |
| KP-B5 | 3,347,985 | T**C**G | Y |
| KP-B6 | 327,129 | T**C**G | Y |
| KP-B6 | 1,400,074 | C**C**G | N |
| KP-B6 | 2,254,942 | T**C**G | Y |
| KP-B6 | 2,318,894 | T**C**G | Y |
| KP-B6 | 3,049,130 | T**C**G | Y |
| KP-B6 | 3,063,632 | C**G**A | Y |
| KP-B6 | 3,133,040 | T**C**C | Y |
| KP-B6 | 3,519,995 | T**C**G | Y |

| | | | |
|---|---|---|---|
| KP-B6 | 4,046,644 | T**C**G | Y |
| KP-B6 | 5,313,678 | T**C**G | N |
| KP-B7 | 1,386,432 | C**G**A | Y |
| KP-B7 | 1,400,074 | C**C**G | N |
| KP-B7 | 3,347,985 | T**C**G | Y |
| KP-B8 | 1,386,432 | C**G**A | Y |
| KP-B8 | 1,400,074 | C**C**G | N |
| KP-B8 | 3,347,985 | T**C**G | Y |

| ***K. pneumoniae* CP000647.1 *rpoB*** | | | |
|---|---|---|---|
| **Isolate** | **Position** | **Context** | **DddA signature (Yes/No)** |
| KP-A1 | 4,771,622 | G**G**A | Y |
| KP-A2 | 4,771,622 | G**G**A | Y |
| KP-A3 | 4,771,622 | G**G**A | Y |
| KP-B1 | 4,771,622 | G**G**A | Y |
| KP-B2 | 4,771,622 | G**G**A | Y |
| KP-B3 | 4,771,622 | G**G**A | Y |
| KP-B4 | 4,771,622 | G**G**A | Y |
| KP-B5 | 4,771,622 | G**G**A | Y |
| KP-B6 | 4,771,622 | G**G**A | Y |
| KP-B7 | 4,771,622 | G**G**A | Y |
| KP-B8 | 4,771,622 | G**G**A | Y |
| KP-C1 | 4,771,622 | G**G**A | Y |
| KP-C2 | 4,771,622 | G**G**A | Y |
| KP-C3 | 4,771,622 | G**G**A | Y |
| KP-C4 | 4,771,622 | G**G**A | Y |
| KP-C5 | 4,771,622 | G**G**A | Y |
| KP-C6 | 4,771,622 | G**G**A | Y |
| KP-C7 | 4,771,622 | G**G**A | Y |
| KP-C8 | 4,771,622 | G**G**A | Y |
| KP-C9 | 4,771,622 | G**G**A | Y |

| ***P. aeruginosa* PAO1 NC_002516.2** | | | |
|---|---|---|---|
| **Isolate** | **Position** | **Context** | **DddA signature (Yes/No)** |
| PAO1-1 | 4,779,056 | G**A**C | N |
| PAO1-3 | 4,779,056 | G**A**C | N |
| PAO1-5 | 4,779,057 | G**G**A | Y |
| PAO1-6 | 4,779,056 | G**A**C | N |
| PAO1-7 | 4,779,056 | G**A**C | N |
| PAO1-8 | 4,779,056 | G**A**C | N |
| PAO1-13 | 4,778,882 | T**C**C | Y |
| PAO1-14 | 4,778,882 | T**C**C | Y |
| PAO1-14 | 4,779,056 | G**A**C | N |
| PAO1-15 | 4,779,057 | G**G**A | Y |
| PAO1-19 | 4,779,057 | G**G**A | Y |

| | | | |
|---|---|---|---|
| PAO1-20 | 4,779,056 | G**A**C | N |
| PAO1-21 | 4,779,011 | T**C**C | Y |
| PAO1-27 | 4,779,056 | G**A**C | N |
| PAO1-31 | 4,779,056 | G**A**C | N |
| PAO1-33 | 4,779,056 | G**A**C | N |
| PAO1-35 | 4,780,163 | C**A**G | N |
| PAO1-38 | 4,779,057 | G**G**A | Y |
| PAO1-39 | 4,779,056 | G**A**C | N |
| PAO1-41 | 4,779,056 | G**A**C | N |
| PAO1-43 | 4,780,163 | C**A**G | N |
| PAO1-43 | 4,779,056 | G**A**C | N |
| PAO1-45 | 4,780,163 | C**A**G | N |
| PAO1-48 | 4,779,056 | G**A**C | N |
| PAO1-50 | 4,779,056 | G**A**C | N |
| PAO1-52 | 4,779,056 | G**A**C | N |
| PAO1-54 | 4,779,056 | G**A**C | N |
| PAO1-56 | 4,779,056 | G**A**C | N |
| PAO1-57 | 4,779,056 | G**A**C | N |
| PAO1-60 | 4,779,056 | G**A**C | N |
| PAO1-61 | 4,779,056 | G**A**C | N |
| PAO1-62 | 4,779,056 | G**A**C | N |
| PAO1-66 | 4,779,056 | G**A**C | N |
| PAO1-67 | 4,779,056 | G**A**C | N |
| PAO1-68 | 4,780,163 | C**A**G | N |
| PAO1-69 | 4,779,056 | G**A**C | N |
| PAO1-70 | 4,779,056 | G**A**C | N |
| PAO1-71 | 4,779,056 | G**A**C | N |
| PAO1-72 | 4,779,056 | G**A**C | N |
| PAO1-74 | 4,779,056 | G**A**C | N |

**Supp. Table 2. X-ray data collection and refinement statistics. Related to Figure 6.**

| | ***P. syringae*** **SsdA$_{tox}$ / SsdA$_{I}$ SAD peak[a]** |
|---|---|
| PDB accession code | 7JTU |
| **Data Collection** | |
| Space group | $I4_122$ |
| Cell dimension | |
| *a, b, c* (Å) | 93.07, 93.07, 383.49 |
| α, β, γ (°) | 90, 90, 90 |
| Wavelength (Å) | 0.9790 |
| Resolution (Å) | 33.4 - 3.0 (3.15-3.0)[b] |
| No. unique reflections | 17597 (434) |
| $R_{merge}$ | 0.12 |
| I/σI | 22.9 (2.7) |
| Completeness (%) | |
| Total | 100 (100) |
| Anomalous | 100 (100) |
| Redundancy | 28.6 (29.6) |
| Wilson B-factor (Å$^2$) | 79.6 |
| **Refinement** | |
| Resolution (Å) | 33.4 – 3.0 (3.1 – 3.0) |
| No. reflections | 17248 (1193) |
| Rwork/Rfree (%) | 20.2 / 23.5 (32.3 / 40.5) |
| No. atoms | |
| Protein | 2717 |
| Ligand/ion | 0 |
| Water | 3 |
| B-factors (Å$^2$) | |
| Protein | 79.6 |
| Ligand/ion | N/A |
| Water | 63.9 |
| rmsd | |
| Bond lengths (Å) | 0.009 |
| Bond angles (°) | 1.086 |
| Missing residues | |
| chain A | 245-257, 409 |
| chain B | none |

[a]All data collected from a single crystal
[b]Values in parentheses are for the highest resolution shell

**Supp. Table 3. Strains, plasmids and primers used in this study.**

| **Bacterial species and strains** | **Genotype** | **Reference/Source** |
|---|---|---|
| *Escherichia coli* DH5α | F– φ80*lacZ*ΔM15 Δ*(lacZYA-argF)*U169 *recA*1 *endA*1 *hsdR*17(rK–, mK+) *phoA* supE44 λ– thi-1 *gyrA*96 *relA*1 | Thermo Fisher Scientific Cat#18258012 |
| *Escherichia coli* BL21 | $F^-$ *omp*T *hsd*$S_B$ ($r_B^-$, $m_B^-$) *gal dcm* (DE3) | EMD Millipore Cat#69450 |
| *Escherichia coli* BL21 *ung-151* | $F^-$ *omp*T *hsd*$S_B$ ($r_B^-$, $m_B^-$) *gal dcm ung-151* | (84) |
| *Escherichia coli* XK1502 | F− Δ*lacU169 nalA* | (85) |
| *Escherichia coli* XK1502 *Δung* | F− Δ*lacU169 nalA* Δung | (28) |
| *Escherichia coli* AB1157 | F- thr-1 leuB6(Am) *glnX*44(AS) *hisG*4(Oc) *rfbC*1 *rpsL*31(strR) *argE*3(Oc) | (86) |
| *Escherichia coli* AB1157 *YPet-dnaN* | *YPet-dnaN,* KanR | (78) |
| *Escherichia coli* CJ236 | $F^+$ *ung-1 relA1 dut-1 spoT1 thiE1* | (87) |
| *Burkholderia cenocepacia* H111 | Wild-type | (88) |
| *Burkholderia cenocepacia* H111 Δ*icmF1* | ΔI35_RS01770 | (28) |
| *Burkholderia cenocepacia* H111 *dddA*$^{E1347A}$ | *dddA*$^{E1347A}$ | (28) |
| *Pseudomonas aeruginosa* PAO1 | Wild-type | (89) |
| *Escherichia coli* O157:H7 EDL933 | Wild-type | (90) |
| *Klebsiella pneumoniae* MGH 78578 | Wild-type | (91) |
| *Acinetobacter baumannii* ATCC 17978 | Wild-type | (92) |
| *Burkholderia thailandensis* E264 | Wild-type | (93) |
| *Pseudomonas putida* F1 | Wild-type | (94) |
| **Plasmids** | **Purpose** | **Reference/Source** |
| pPSV39-CV | For inducible expression of proteins in *E. coli* | (95) |

| | | |
|---|---|---|
| pScrhaB2-V | For inducible expression of proteins in *E. coli* | (96) |
| pEXG2 | For generation of markless *P. aeruginosa* mutants | (74) |
| pScrhaB2-V::*ssdA* | To express *ssdA* | This study |
| pPSV39-CV::$ssdA_I$ | To express *ssdAI* | This study |
| pScrhaB2-V::*TequE* | To express *ssdA* | This study |
| pPSV39-CV::*TequE* | To express *ssdAI* | This study |
| pETDuet-1 mcs1::$ssdA\text{-}his_6$ $mcs1::ssdA_I$ | To co-express $dddA\text{-}dddA_I$ | This study |
| pexG2_Δ*ung* | To delete Δ*ung* in *P. aeruginosa* PAO1 | This study |
| **Primers and gBlocks** | **Sequence** | **Purpose** |
| ungDel-1 | 5'-CAAGCTTCTGCAGGTCGACTCTAGAGGTATGGAGTTGTCCTTCGG | To clone ung deletion cassete into pEXG2 |
| ungDel-2 | 5-AGAGGTCCGGATCGGTCATGGAACCCCC | To clone ung deletion cassete into pEXG2 |
| ungDel-3 | 5'-CATGACCGATCCGGACCTCTGAAGGCCGC | To clone ung deletion cassete into pEXG2 |
| ungDel-4 | GGAAATTAATTAAGGTACCGAATTCCCGCGCCGGTGGACTGGC | To clone ung deletion cassete into pEXG2 |
| PAO1-ung-F | 5'-CCGGGGAGTACTTCTCGTTC | To confirm *ung* deletion in P. aeruginosa PAO1 |
| PAO1-ung-R | 5'-GGCGTTCCAGTACCTGCTC | To confirm *ung* deletion in P. aeruginosa PAO1 |
| GA_duet_PsyrE1-F | 5'-ACCATCATCACCACAGCCAGGATCCGAAGGTCTCAAATATTGCG | To clone *ssdA* into MCS1 of pETDuet |
| GA_duet_PsyrE1-R | 5'-CTTAAGCATTATGCGGCCGCTCATTCCGACCTCATAATTG | To clone *ssdA* into MCS1 of pETDuet |
| GA_duet_PsyrI1-F | 5'-TATAAGAAGGAGATATACATATGAATAACAAAAGTAAAGTATTGATTGAAAAGC | To clone $ssdA_I$ into MCS1 of pETDuet |
| GA_duet_PsyrI1-R | 5'-GCCGGCCGATATCCAATTGAGATCTTCACACAACTTGCGGCAC | To clone $ssdA_I$ into MCS1 of pETDuet |
| GA_pRhB_PsyrE1-F | 5'-TGAAATTCAGCAGGATCACATATGAAGGTCTCAAATATTGCG | To clone *ssdA* into pSCRhaB2 |

| | | |
|---|---|---|
| GA_pRhB_PsyrE1-R | 5’-TCATTTCAATATCTGTATATCTAGATTCCGACCTCATAATTGTTTC | To clone ssdA into pSCRhaB2 |
| GA_p39_PsyrI1-F | 5’-ACAATTTCAGAATTCGAGCTCACGGGAGGAAAGATGAATAACAAAAGTAAAGTATTGATTGAAAAGC | To clone *ssdA*$_I$ into pPSV39 |
| GA_p39_PsyrI1-R | 5’-TCATTTCAATATCTGTATATCTAGATCACACAACTTGCGGCAC | To clone *ssdA*$_I$ into pPSV39 |
| rpoB-F | 5’-GGAAAACCAGTTCCGCGTTG | To sequence rpoB in Enterobacteriaceae |
| rpoB-R | 5’-TCCAAGTTGGAGTTCGCCTG | To sequence rpoB in Enterobacteriaceae |
| nfo_del-F | 5’-CATTACCGTTTTCCTCCAGCGGGTTTAACAGGAGTCCTCGCATGAAATACGTGTAGGCTGGAGCTGCTTC | To generate *nfo* deletion cassete |
| nfo_del-R | 5’-CCGTAAAATTGCAAGGATCTCCTTTTCCCGGTTATTCATCTTCAGGCTACCATATGAATATCCTCCTTAG | To generate *nfo* deletion cassete |
| nfo_dt-F | 5’-GCTGATGGCACTGGTACTGT | To confirm *nfo* deletion |
| nfo_dt-R | 5’-CCTTTAATCCGGCCTTTGCG | To confirm *nfo* deletion |
| xthA_del-F | 5’-TACCATCCACGCACTCTTTATCTGAATAAATGGCAGCGACTATGAAATTTGTGTAGGCTGGAGCTGCTTC | To generate *xthA* deletion cassete |
| xthA_del-R | 5’-TTAATTCTCCTGACCCAGTTTGAGCCAGGAGAGCTGCTAAATTAGCGGCGCATATGAATATCCTCCTTAG | To generate *xthA* deletion cassete |
| xthA_dt-F | 5’-TACGTTTGCGATGTGGGTGA | To confirm *xthA* deletion |
| xthA_dt-R | 5’-ATAACAAAGGACGGCAGGCA | To confirm *xthA* deletion |

| | | |
|---|---|---|
| EL142_RS06975[tox]-gBlock | 5’-<br>TGAAATTCAGCAGGATCACATATGCTT<br>TTAGGTGGACTTAACAACTACCAATAC<br>GCCCCAAATCCAGTCGAATGGGTCGAT<br>CCTTTGGGTTGGAAATTCTCCAATGGC<br>AAGCGTCGTCCGCCCCACAAGGCAAC<br>GGTTACCGTCACAGACAAGAACGGAG<br>TGGTCAAACACAAATCCAATTTGGTGT<br>CAGGAAATATGACAGAAGCCGAAAAA<br>AAACTGGGTTTCCCGAACAACTCTTTG<br>GCAACACATACCGAGAATCGTGCAAC<br>GCGCTTAATTGACCTGAATCAAGGTGA<br>TACCATGTTAATTGAAGGACAGTATCG<br>CCCGTGCCCACGCTGTAAGGGTGCGAT<br>GCGTGTTAAGGCAGAGGAATCTGGGG<br>CTAAGGTTATCTACACCTGGCCCGAAG<br>ACGGTGACTTGAAGAAGCGCGAGTGG<br>GAAGGAACCCCCTGTGATAAAAAGTC<br>TAGATATACAGATATTGAAATGA | gBlock to clone TequE into pSChRhaB2 |
| EL142_RS06970 - gBlock | 5’-<br>GCCCCAAGGGGTTATGCTAAAGCTTTC<br>TTAATTATCGAACAGGAATTTCAGATG<br>ATTCTCGACTTCAAGAATGTCAATCTT<br>TTCAATCAAGCGAAGACGAACGTAGTT<br>ATTAGGCTCAATTAAAAGATCGACACA<br>ATTTGGTTCGGACAGTAAATAGCCGGA<br>CAAGCCATAGGGGTTAAAATCAAGGA<br>ACAGATTAGACTCCAGATAGATGTCTC<br>CCTCGGGGATACCACCGCTGCGTTCCG<br>TTGTTGAACGTGTAATGATGAAATACT<br>GCTCAGGCACTTCATAGCTATCAGATG<br>CAAACGAAAGGATGGCGCAATAATCC<br>TCAATCGCAAACTTCACTTCTTGAATC<br>ACGATGTTACTCAGCATCTTTCCTCCC<br>GTGAGCTCGAATTCTGAAATTGT | gBlock to clone TequE into pPSV39 |
| **Substrates and oligonucleotides for biochemical analysis** | **Sequence** | **Purpose** |
| RNA-GGCCGG | 5’-<br>GAGGCCGGAAGUGGAUGUGGAUAAG<br>AUGGAG | RNA substrate for deaminase biochemical reaction |
| RNA-GGCUGG | 5’-<br>GAGGCUGGAAGUGGAUGUGGAUAAG<br>AUGGAG | RNA substrate for deaminase biochemical reaction |

| | | |
|---|---|---|
| PPE-Oligonucleotide | 5’FAM-CTCCATCTTATCCACATCCACT | Oligonucleotide for poisoned primer extension assay |
| DNA-ATGCGCCA | 5’FAM-AAAAAAAAAAAAAA<br>ATGCGCCAAAAAAAAAAAAAAA | DNA substrate for deaminase biochemical reaction |
| revDNA-ATGCGCCA | 5’-<br>TTTTTTTTTTTTTTTGGCGCATTTTTTTT<br>TTTTTTT | Reverse complement to generate dsDNA substrate with DNA-ATGCGCCA |
| DNA-GCG | 5’-FAM-AAAAAAAAAAAAAAAA<br>GCGAAAAAAAAAAAAAAAAA | DNA substrate for deaminase biochemical reaction |
| DNA-CCG | 5’-FAM-AAAAAAAAAAAAAAAA<br>CCGAAAAAAAAAAAAAAAAA | DNA substrate for deaminase biochemical reaction |
| DNA-TCG | 5’-FAM-AAAAAAAAAAAAAAAA<br>TCGAAAAAAAAAAAAAAAAA | DNA substrate for deaminase biochemical reaction |
| DNA-ACG | 5’-FAM-AAAAAAAAAAAAAAAA<br>ACGAAAAAAAAAAAAAAAAA | DNA substrate for deaminase biochemical reaction |
| revDNA-GCG | 5’-<br>TTTTTTTTTTTTTTTTTCGCTTTTTTTTTT<br>TTTTTT | Reverse complement to generate dsDNA substrate with DNA-CGC |
| revDNA-CCG | 5’-<br>TTTTTTTTTTTTTTTTTCGGTTTTTTTTT<br>TTTTTTT | Reverse complement to generate dsDNA substrate with DNA-CCG |
| revDNA-TCG | 5’-<br>TTTTTTTTTTTTTTTTTCGATTTTTTTTT<br>TTTTTTT | Reverse complement to generate dsDNA substrate with DNA-TCG |
| revDNA-ACG | 5’-<br>TTTTTTTTTTTTTTTTTCGTTTTTTTTTT<br>TTTTTT | Reverse complement to generate dsDNA substrate with DNA-ACG |
